# Mechanistic Insights into the *Cis-* and *Trans*-acting Deoxyribonuclease Activities of Cas12a

**DOI:** 10.1101/353748

**Authors:** Daan C. Swarts, Martin Jinek

## Abstract

**HIGHLIGHTS:**

- Target ssDNA binding allosterically induces unblocking of the RuvC active site
- PAM binding facilitates unwinding of dsDNA targets
- Non-target DNA strand cleavage is prerequisite for target DNA strand cleavage
- After DNA cleavage, Cas12a releases the PAM-distal DNA product

**SUMMARY:** CRISPR-Cas12a (Cpf1) is an RNA-guided DNA-cutting nuclease that has been repurposed for genome editing. Upon target DNA binding, Cas12a cleaves both the target DNA in *cis* and non-target single stranded DNAs (ssDNA) in *trans.* To elucidate the molecular basis for both deoxyribonuclease cleavage modes, we performed structural and biochemical studies on *Francisella novicida* Cas12a. We show how crRNA-target DNA strand hybridization conformationally activates Cas12a, triggering its *trans*-acting, non-specific, single-stranded deoxyribonuclease activity. In turn, *cis*-cleavage of double-stranded DNA targets is a result of PAM-dependent DNA duplex unwinding and ordered sequential cleavage of the non-target and target DNA strands. Cas12a releases the PAM-distal DNA cleavage product and remains bound to the PAM-proximal DNA cleavage product in a catalytically competent, *trans*-active state. Together, these results provide a revised model for the molecular mechanism of Cas12a enzymes that explains their *cis*- and *trans*-acting deoxyribonuclease activities, and additionally contribute to improving Cas12a-based genome editing.

## INTRODUCTION

In prokaryotes, CRISPR-Cas systems (clustered regularly interspaced palindromic repeats and CRISPR-associated proteins) function as programmable immune systems that utilize CRISPR RNAs (crRNAs) as guide molecules for the recognition and targeting of invasive nucleic acids such as viral DNA (Koonin et al., 2017; Mohanraju et al., 2016). Class 2 CRISPR-Cas systems, which include type II and V systems, elicit antiviral immunity by means of single multidomain nuclease enzymes that mediate crRNA-guided target DNA binding and catalyze target DNA cleavage. Cas9 and Cas12a proteins, the respective effector nucleases of type II and type V-A systems, mediate crRNA-guided DNA cleavage by recognizing two distinct sequence elements in the target DNA: (i) a ‘protospacer’ sequence, specified by Watson-Crick base-pairing interactions with the crRNA, and (ii) a protospacer adjacent motif (PAM), recognized via sequence- and/or shape-specific protein-DNA interactions (Swarts and Jinek, 2018). Upon binding of a cognate target DNA, the complementary spacer-derived segment of the crRNA and the protospacer ‘target strand’ (TS) form an RNA-DNA heteroduplex. At the same time, the non-target strand (NTS) of the protospacer is displaced. Formation of this so-called ‘R-loop’ structure catalytically activates both Cas9 and Cas12a, triggering *cis*-cleavage of both target DNA strands. The ability to program Cas9 and Cas12a with a crRNA sequence of choice has led to their repurposing for precision genome editing and for regulation of gene expression (Barrangou and Doudna, 2016; Doudna et al., 2016; Swarts and Jinek, 2018; Wang et al., 2016).

Despite similar functionalities, Cas9 and Cas12a proteins have distinct evolutionary histories and therefore distinct mechanistic properties (Shmakov et al., 2017; Swarts and Jinek, 2018). *Streptococcus pyogenes* Cas9, which is widely used for genome editing, recognizes a 5’- NGG-3’ PAM directly downstream of the protospacer NTS (Anders et al., 2014). In contrast, Cas12a orthologues used for genome editing recognize a 5’-TTTV-3’ PAM directly upstream of the protospacer TS (Gao et al., 2016; Yamano et al., 2016; Zetsche et al., 2015). Cas9 and Cas12a furthermore use different mechanisms for DNA cleavage. Cas9 cleaves the TS and NTS with its HNH and RuvC domains, respectively (Jinek et al., 2012). This typically results in PAM-proximal DNA breaks with blunt-ends or 1-nucleotide overhangs. In contrast, Cas12a uses a single RuvC nuclease domain for cleavage of both strands of the target DNA, generating PAM-distal DNA breaks with larger (5-7 nt) 5’ overhangs (Swarts et al., 2017). Strikingly, Cas12a has two DNA cleavage modes (Chen et al., 2018; Li et al., 2018). Besides canonical crRNA-guided *cis*-cleavage of target DNA, crRNA-TS DNA hybridization triggers sequence-independent deoxyribonuclease activity in Cas12a that cleaves and degrades nontarget single-stranded DNA in *trans*.

Structural studies of Cas12a have revealed its overall molecular architecture and mechanism of target DNA recognition. The enzyme has a bilobed structure in which the N-terminal REC lobe, consisting of the α-helical domains REC1 and REC2, is connected by the Wedge (WED) domain to the C-terminal NUC lobe comprising the PAM interacting (PI), Bridge Helix (BH), RuvC and Nuc dmains (Dong et al., 2016; Gao et al., 2016; Stella et al., 2017; Swarts et al., 2017; Yamano et al., 2016). The crRNA is recognized specifically via shape-specific recognition of the repeat-derived part of the crRNA, which forms a pseudoknot structure (Dong et al., 2016). crRNA binding results in structural pre-ordering of the seed sequence in the crRNA, thereby priming the Cas12-crRNA complex for target DNA recognition (Swarts et al., 2017). Shape- and sequence-specific interactions with the PAM then facilitate target dsDNA binding and is prerequisite for catalyzing double-strand break formation (Gao et al., 2016; Yamano et al., 2016; Zetsche et al., 2015). Despite these insights, the molecular mechanisms underpinning *cis*- and *trans*-acting deoxyribonuclease activities of Cas12a remain largely unclear.

To obtain insights into both *cis*- and *trans*-acting DNA cleavage activities of Cas12a, we have determined the crystal structures of *Francisella novicida* U112 Cas12a (FnCas12a) in ternary complex with a crRNA and either ssDNA or dsDNA targets. Together with corroborating biochemical experiments, the structure of Cas12a bound to a guide crRNA and singlestranded DNA target reveals that crRNA-TS DNA duplex formation, rather than PAM binding, drives a conformational rearrangement required for catalytic activation of the Cas12a RuvC domain. Although PAM binding does not play a role in the catalytic activation of Cas12a, it is essential for the recognition and cleavage of dsDNA targets since it promotes dsDNA strand separation and crRNA-TS DNA hybridization. We have additionally determined a crystal structure of Cas12a-crRNA in complex with a dsDNA target, revealing that the NTS of the target DNA is electrostatically guided toward the catalytic site in the RuvC domain, thereby eliciting sequential cleavage of the NTS and TS DNA strands. We furthermore demonstrate that after target DNA cleavage by Cas12a, the PAM proximal-DNA cleavage product remains bound to the Cas12a-crRNA complex. This maintains Cas12a in a catalytically activated state, allowing for degradation of non-target single stranded DNA in *trans*. In contrast, the PAM distal-DNA cleavage product is released from Cas12a.

## RESULTS

### Target ssDNA binding allosterically activates the RuvC catalytic site in Cas12a

In addition to cleaving target dsDNAs in *cis*, Cas12a enzymes were recently reported to possess *trans* DNA cleavage activity triggered by target ssDNA binding (Chen et al., 2018; Li et al., 2018). To corroborate these findings, we programmed *Francisella novicida* Cas12a (FnCas12a) with a crRNA and incubated the resulting complex with circular ssDNA (M13 ssDNA) or circular dsDNA (M13 dsDNA) substrates that bear no complementarity to the crRNA. Next, we added to the reaction 20-nt ssDNA oligonuclotides that lacked a PAM and were either complementary or lacked complementarity to the spacer-derived segment of the crRNA (**Figure 1A**). In the presence of a non-complementary ssDNA, FnCas12a did not degrade the M13 DNAs. In contrast, the presence of a complementary target ssDNA caused complete degradation of M13 ssDNA, but not the M13 dsDNA. This activity was not observed for the RuvC catalytic site mutant FnCas12a^E1006Q^, indicating that the observed in *trans* degradation of ssDNA substrates is mediated by the RuvC catalytic site that also mediates canonical in *cis* cleavage of target dsDNA. These findings are in agreement with the recently reported *trans*-acting activity of Cas12a (Chen et al., 2018; Li et al., 2018) and suggest that crRNA-TS hybridization alone drives allosteric activation of Cas12a.

**Figure 1.**
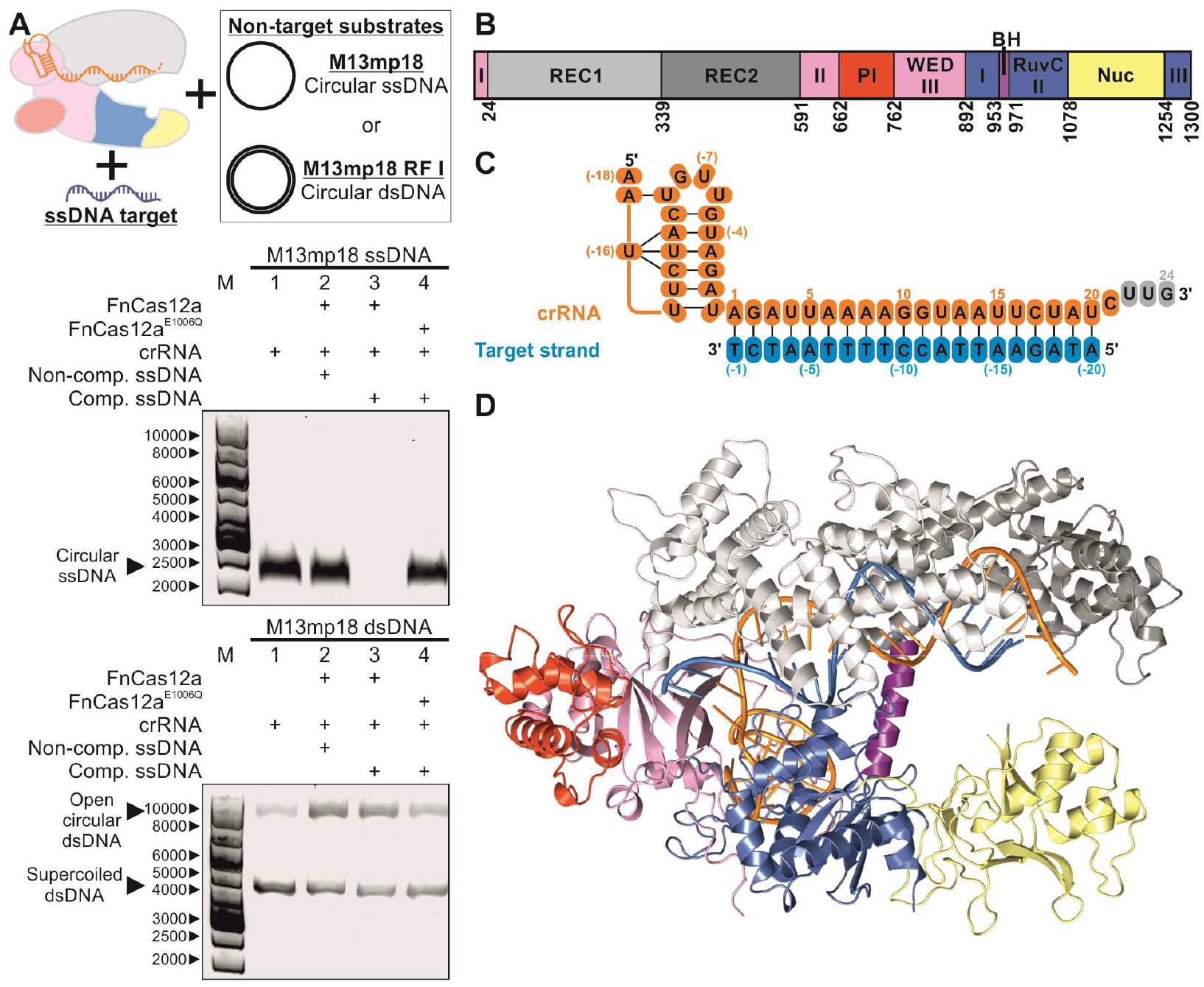
Structure of FnCas12a-crRNA in complex with a ssDNA Target. (A) A single stranded target DNA activates unspecific ssDNase activity of FnCas12a. Top: schematic representation of the experiment. Middle and Bottom: FnCas12a-crRNA complexes were incubated with or without (non-)complementary ssDNA target and non-target circular ssDNA (middle) or dsDNA (bottom) substrates. Degradation products were resolved by agarose gel electrophoresis. M: 1kb DNA ladder marker, Comp.: complementary. (B) Schematic diagram of the domain organization of FnCas12a. REC, recognition; PI, protospacer adjacent motif (PAM) interacting; WED, wedge; BH, bridge helix; Nuc, nuclease. Both the WED and RuvC domains are formed by three discontinuous segments of the protein sequence. (C) Sequence of the crRNA guide and TS DNA (see also Table S1) in the structure of the FnCas12a-crRNA-TS complex. Structurally disordered nucleotides are colored gray. (D) Overall structure of the FnCas12a-crRNA-TS complex. Domains are colored according to the scheme in panel B.

To obtain structural insights into the allosteric mechanism underpinning catalytic activation of Cas12a, we determined the crystal structure of a ternary complex of FnCas12a bound to a crRNA guide and a 20-nt PAM-less ssDNA TS (**Figure 1B-D, Table 1**) at a resolution of 2.8 Å. In agreement with other Cas12a structures, the FnCas12a-crRNA-TS structure preserves a bilobed architecture. The first 20 nucleotides of the spacer-derived segment of the crRNA (A1-U20) form Watson-Crick base pairs with the TS, while the 3’-terminal nucleotides of the crRNA (U22-G24) are unordered.

**Table 1.** Crystallographic Data Collection and Refinement Statistics

| Dataset | TS only complex | Ternary complex |
| --- | --- | --- |
| FnCas12a | E1006Q | WT |
| RNA | crRNA1 | crRNA1 |
| DNA | oDS311 (TS) | oDS142 (TS) + oDS141 (NTS) |
| X-ray source | SLS PXIII | SLS PXIII |
| Space group | $P2_1 2_1 2_1$ | $P1 2_1 1$ |
| Cell dimensions |  |  |
| a, b, c (Å) | 78.80, 788.58, 284.12 | 82.17, 141.68, 89.48 |
| $\alpha$, $\beta$, $\gamma$ (°) | 90, 90, 90 | 90, 97.43, 90 |
| Wavelength (Å) | 1.000 | 1.008 |
| Resolution (Å)* | 49.14–2.80 (2.9–2.8) | 47.62–2.65 (2.745–2.65) |
| $R_{\text{merge}}$ (%)* | 0.21 (3.234) | 0.075 (1.761) |
| CC1/2 (%)* | 99.9 (33.3) | 100 (63.8) |
| $I/\sigma I$* | 16.15 (1.20) | 17.86 (0.96) |
| Completeness (%)* | 99.96 (99.99) | 99.59 (98.67) |
| Redundancy* | 26.1 (24.9) | 6.9 (6.8) |
| Refinement |  |  |
| Resolution (Å) | 2.80 | 2.65 |
| No. reflections | 105,119 | 58,701 |
| $R_{\text{work}} / R_{\text{free}}$ | 0.212/0.240 | 0.246/0.261 |
| No. atoms |  |  |
| Protein | 20893 | 10578 |
| Nucleic acid | 2476 | 2265 |
| Ion/ligand | 207 | 108 |
| Water | 52 | 30 |
| B Factors |  |  |
| Mean (Å<sup>2</sup>) | 87.42 | 106.99 |
| Protein (Å<sup>2</sup>) | 79.13 | 105.19 |
| Nucleic acid (Å<sup>2</sup>) | 64.49 | 116.62 |
| Ion/ligand (Å<sup>2</sup>) | 99.54 | 88.23 |
| Water (Å<sup>2</sup>) | 67.70 | 78.86 |
| RMSDs |  |  |
| Bond lengths (Å) | 0.005 | 0.004 |
| Bond angles (°) | 0.93 | 0.91 |
| Ramachandran plot |  |  |
| Favoured (%) | 96.80 | 97.96 |
| Allowed (%) | 3.16 | 2.04 |
| Outliers (%) | 0.04 | 0.00 |
| Molprobrity |  |  |
| Clashscore | 6.83 | 8.26 |
| * Values in parentheses denote highest resolution shell |  |  |

Previously determined structures of binary Cas12a-crRNA complexes (PDB: 5NG6) showed that the REC2 domain occludes the catalytic site in the RuvC domain (**Figure 2A**), which likely prevents not only in *cis* target DNA cleavage, but also in *trans* ssDNA degradation. This is consistent with the observed lack of *cis*- or *trans*-nuclease activity in the FnCas12a-crRNA complex in the absence of a cognate target DNA (**Figure 1A**). Structural superposition of the binary FnCas12a-crRNA (PDB: 5NG6) and ternary FnCas12a-crRNA-TS complex structures reveals the conformational changes associated with catalytic activation by TS binding (**Figure 2A**; root-mean-square deviation (RMSD): 12.54 Å). The NUC lobe is largely unperturbed by TS binding (RMSD of NUC lobe alignment: 2.61 Å), with only a small rotation of the Nuc domain by ~9° relative to the remainder of the NUC lobe. In contrast, the REC lobe undergoes a major conformational rearrangement (RMSD of REC lobe alignment: 15.26 Å). The REC1 domain rotates approximately 33° relative to the NUC lobe and thereby moves closer to the NUC lobe and to the crRNA-TS heteroduplex, allowing formation of stabilizing interactions between the crRNA-TS heteroduplex and REC1 (**Figure S1**). The REC2 domain undergoes a rotation of 46° and a translation of ~9.5 Å relative to the REC1 domain. This is essential to accommodate the PAM-distal end of the crRNA-TS heteroduplex, which would otherwise result in severe clashes with the REC2 domain (**Figure S2**). In addition, the rearrangement results in the formation of stabilizing interactions between the crRNA-TS heteroduplex and REC2 residues (**Figure S1B**). The conformational changes induced by progressive crRNA-TS DNA hybridization thus move the REC2 domain away from the NUC lobe, thereby exposing the catalytic site in the RuvC domain and permitting substrate DNA access (**Figure 2A**).

**Figure 2.**
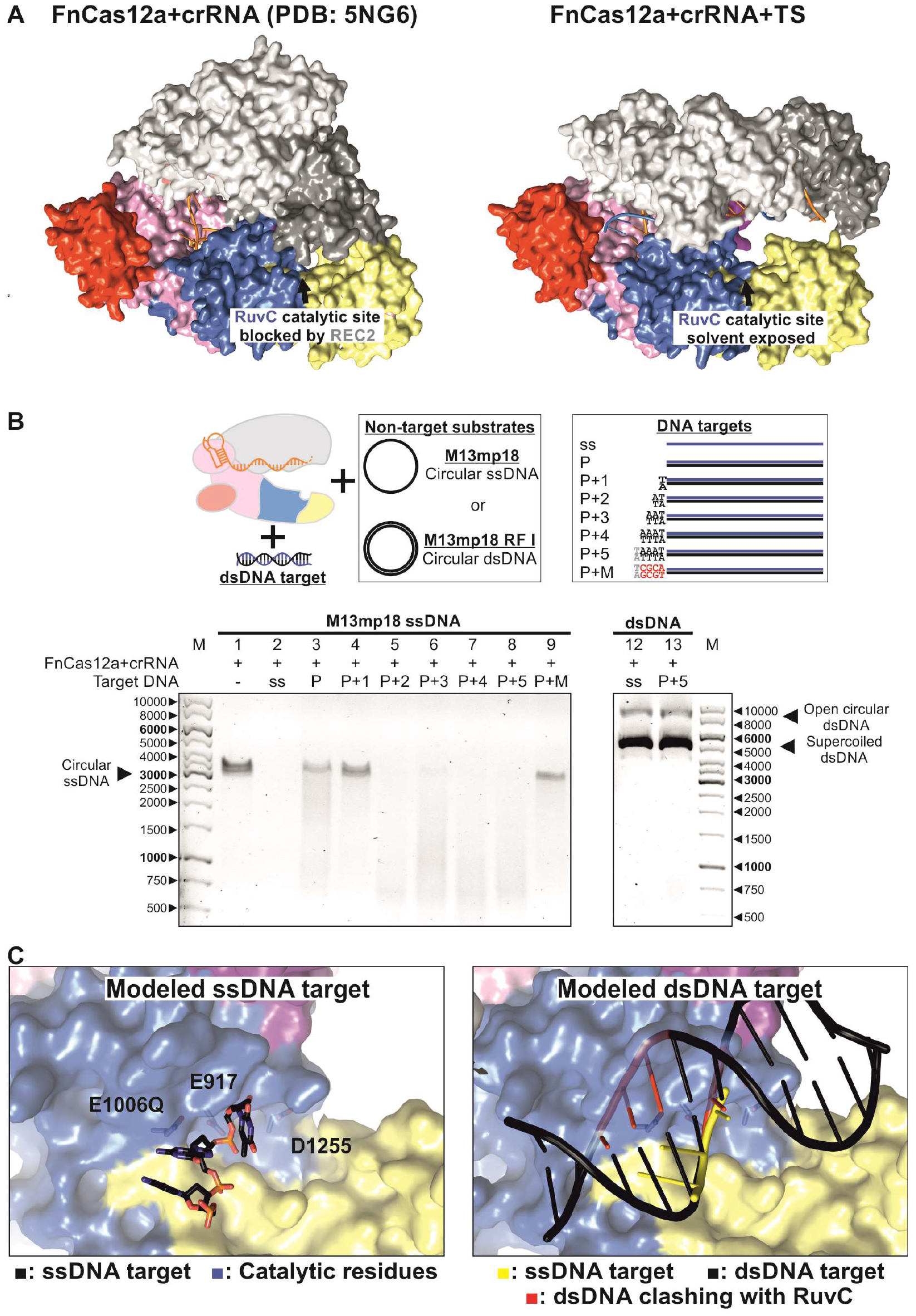
Conformational activation of the FnCas12a RuvC catalytic site. (A) Surface representation of the binary FnCas12a-crRNA (PDB: 5NG6) and the ternary FnCas12a-crRNA-TS complex structures. Domains are colored according to the scheme in Figure 1 panel B. (B) Target dsDNA activates FnCas12a *trans*-acting ssDNA degradation only when it contains a PAM. Top left: schematic representation of the experiment. Top right: schematic representation of the trans-activating DNA targets. Bottom: FnCas12a-crRNA complexes were incubated with various dsDNA targets and non-target circular ssDNA substrates. Degradation products were resolved by agarose gel electrophoresis. M: 1kb DNA ladder marker. Target DNA sequences are provided in Table S1. (C)DNA substrates modeled into the activated catalytic site of FnCas12a. Left: A single stranded target DNA modeled in the RuvC catalytic site of the FnCas12a (PDB: 5NFV). The modeled DNA is based on the structure of AacCas12b-crRNA bound to a DNA target (PDB: 5U33). Right: A double stranded DNA (PDB: 1BNA) modeled in the RuvC catalytic site of the FnCas12a (PDB: 5NFV).

### PAM recognition is essential for target dsDNA unwinding

The observation that target ssDNA binding is sufficient for allosteric activation of Cas12a implies that neither PAM recognition nor NTS coordination are per se required for the allosteric activation of the RuvC nuclease domain. While it has previously been demonstrated that PAM-containing double-stranded DNA targets also allosterically induces the *trans*-acting ssDNase activity of Cas12a (Chen et al., 2018), the significance of the PAM for dsDNA-induced *trans*-nuclease activity remains unclear. To investigate this, we extended our *trans*-acting nuclease activity assays to include complementary dsDNA targets containing canonical, mutated, and truncated PAM sequences (**Figure 2B**). Double-stranded DNA targets containing one or more T nucleotides of the canonical 5’-TTTV-3’ PAM triggered efficient M13 ssDNA degradation. In contrast, almost no M13 ssDNA degradation was observed with dsDNA targets with truncated PAM sequences. Thus, dsDNA-induced *trans*-acting ssDNAse activity requires the presence of at least a 5’-TV-3’PAM in the dsDNA target. A plausible explanation for these observations is that PAM binding is required to initiate unwinding of the dsDNA target and thereby facilitates base pairing between the TS and the seed segment of the crRNA, which is followed by progressive crRNA-TS hybridization, which results in allosteric activation of Cas12a.

Furthermore, although ssDNA and dsDNA target binding induced degradation of the M13 ssDNA substrate in *trans*, neither target DNA was able to trigger degradation of M13 dsDNA substrates in *trans* (**Figure 1A**, **Figure 2B**). We previously modelled the binding of a ssDNA substrate in the RuvC catalytic site based on superpositions with the structures DNA-bound complexes of Cas12b (Swarts et al., 2017). In agreement with the *trans*-cleavage experiments, modeling the binding of a dsDNA substrate indicates that the RuvC dsDNA substrate cannot accommodate a B-form DNA duplex (PDB: 1BNA) due to severe steric clashes of both dsDNA strands with the RuvC domain (**Figure 2C**). This implies that *cis*-cleavage of a dsDNA targets requires that the dsDNA be unwound to allow sequential cleavage of TS and NTS strands in the RuvC active site.

### Surface electrostatics of the REC lobe orchestrates NTS cleavage

Previously determined structures of FnCas12a complexes with dsDNA targets in precleavage (PDB: 5NFV) and post-cleavage (PDB: 5MGA) states revealed that guide crRNA-TS hybridization displaces 20 nucleotides in the NTS (Stella et al., 2017; Swarts et al., 2017). As the displaced segment of the NTS contains the cleavage site, crRNA-TS hybridization would thus make the NTS susceptible to the ssDNase activity of the RuvC domain. However, there is currently little insight into the mechanism coupling crRNA-TS DNA hybridization to NTS DNA cleavage since an extensive portion of the NTS is structurally disordered in these complexes. To shed light on the mechanism of crRNA-guided target DNA cleavage, we determined an additional structure of a ternary complex comprising wild-type FnCas12a, a 43-nt crRNA, and a target dsDNA comprising full-length target and non-target strands. Although the complex contains the same dsDNA ligand as our previously reported precleavage structure (PDB: 5NFV), the new crystal structure has significantly different unit cell dimensions, indicative of a slightly different conformational state (RMSD: 1.25). The structure, determined at 2.65 Å resolution, nevertheless still corresponds to a pre-cleavage state containing a near-complete R-loop in which NTS nucleotides 1-13 and 19-20 are ordered. NTS nucleotides 7-13 are bound in a positively charged groove on the outer surface of the RuvC domain by electrostatic contacts with Asn1288 and basic residues including Lys895, Arg1014, Lys1066, Lys1069, Lys1281, and Lys1287 (**Figure 3A-C, Figure S3**). In part, these interactions likely contribute to energetic stabilization of the displaced NTS in the context of the R-loop, thereby promoting crRNA-TS DNA hybridization. Additionally, the NTS binding groove provides a positively charged conduit connecting the PAM interaction domain and the RuvC active site and thus likely positions the NTS for insertion into the RuvC catalytic site (**Figure 3C, Figure S3**). The positively charged residues involved in NTS binding are partially conserved across the Cas12a nuclease family and the positively charged NTS binding groove is a prominent feature in other Cas12as orthologs (**Figure S4, S5**).

**Figure 3.**
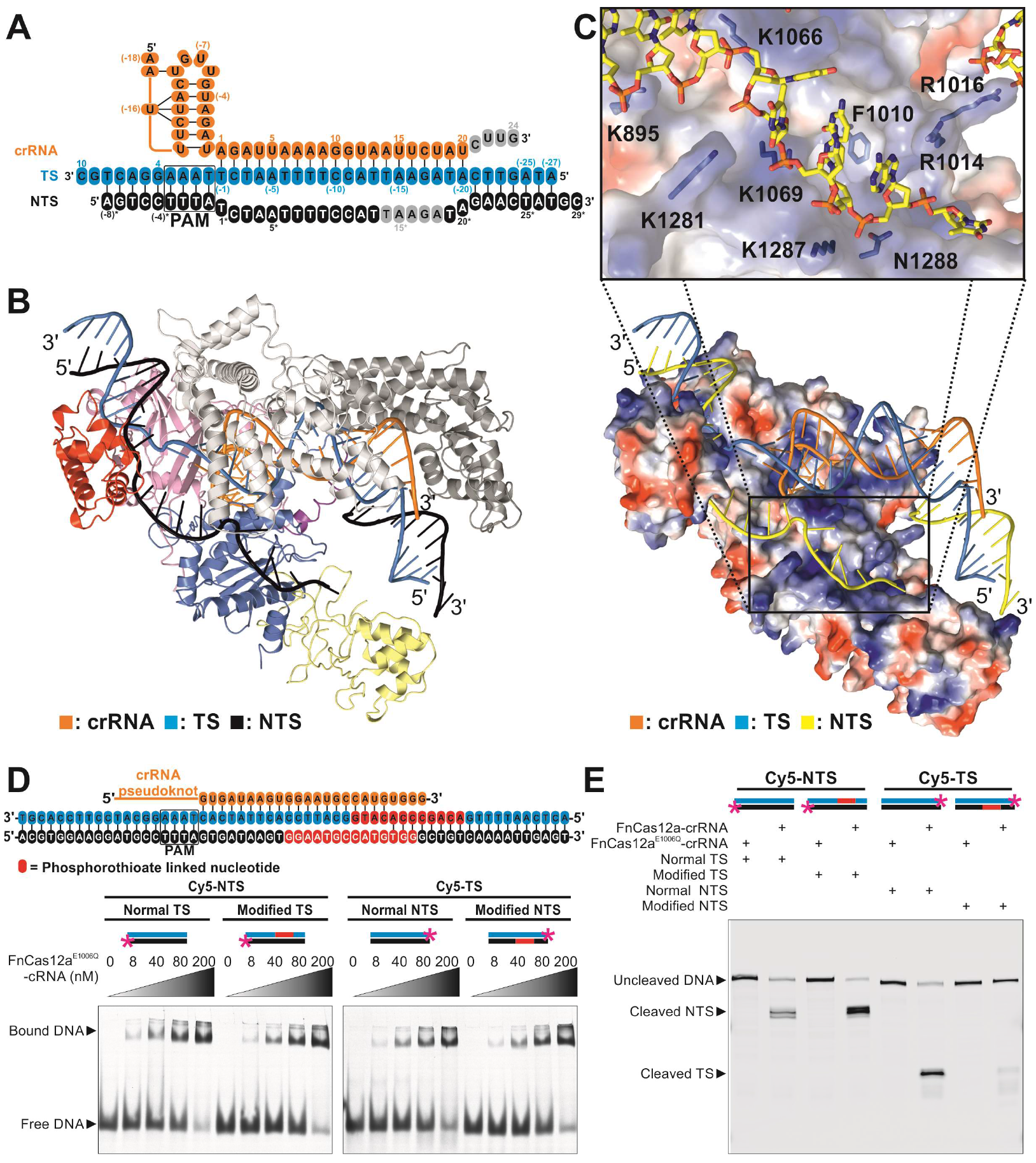
Non-target DNA strand cleavage is prerequisite for target DNA strand cleavage. (A) Sequences of the crRNA guide and TS and NTS DNA (see also Table S1) in the structure of the FnCas12a-crRNA-dsDNA complex. Structurally disordered nucleotides are colored gray. (B) Overall structure of the FnCas12a-crRNA-dsDNA complex. Domains are colored according to the scheme in Figure 1B. (C) Surface electrostatic potential map of the FnCas12a NUC lobe reveals the NTS binding groove. The REC lobe is omitted for clarity. Blue, positively charged region; red, negatively charged region. The inset panel displays the positively charged residues involved in NTS coordination. (D) Phosphorothioate-modification of the DNA backbone does not affect dsDNA binding by Cas12a. Top: sequences of TS and NTS DNA used for the experiments in panel D and E. Phosphorothioate nucleotides in the modified TS and NTS DNAs are colored red. Bottom: FnCas12a^E1006Q^-crRNA complexes were incubated with dsDNA targets and complexes were resolved by native 8% polyacrylamide gel electrophoresis. Bottom left: EMSA using DNAs consisting of a Cy5-labeled NTS and normal or phosphorothioate-modified TS DNAs. Bottom right: EMSA using DNAs consisting of a Cy5-labeled TS and normal or phosphorothioate-modified NTS DNAs. (E)Cas12a cleaves the NTS and TS sequentially. FnCas12a-crRNA complexes were incubated with dsDNA targets and cleavage products were resolved by denaturing (7 M Urea) 20% polyacrylamide gel electrophoresis. While phosporothioate modification of the TS does not affect cleavage of the Cy5-labeled NTS, phosphorothioate modification of the NTS inhibits cleavage of the Cy5-labeled NTS.

### NTS DNA cleavage is prerequisite for TS cleavage

In our previous study, we provided evidence suggesting that Cas12 family enzymes generate double-strand DNA breaks by sequential cleavage of the two substrate DNA strands (Swarts et al., 2017). Based on the observation that the NTS binding groove in FnCas12a appears to guide the unpaired NTS into the RuvC catalytic site (**Figure 3C**), we hypothesized that during dsDNA cleavage the NTS is cleaved before the TS. To test this, we designed modified target dsDNA substrates in which either the TS or the NTS contained multiple backbone phosphorothioate modifications (**Figure 3D**). Such modifications typically inhibit cleavage by divalent cation dependent nucleases (Eckstein and Gish, 1989). The phosphorothioate modifications did not affect FnCas12a-crRNA binding to the dsDNA targets since FnCas12a^E1006Q^-crRNA bound both TS-modified and NTS-modified dsDNA targets equally well as to unmodified control DNAs (**Figure 3D**). Upon incubation with a catalytically active FnCas12a-crRNA complex, we observed that phosphorothioate modification of the TS did not affect NTS cleavage (**Figure 3E**). This indicates that the NTS cleavage can occur independently of TS cleavage. In contrast, TS cleavage was almost completely abrogated in the dsDNA substrate in which the NTS was modified, pointing to its dependence on prior NTS cleavage (**Figure 3E**). One possible explanation is that the presence of an intact NTS in a dsDNA substrate sterically hinders the binding of the TS in the RuvC catalytic site. Alternatively, NTS cleavage is required for the unwinding of the PAM-distal dsDNA segment, which has to occur before the TS can enter the catalytic site. Irrespective of the actual cause of the observed inhibition of TS cleavage, these observations imply an ordered sequential mechanism of dsDNA cleavage in which the NTS is cleaved first and the TS second.

### Cas12a releases the PAM-distal end of cleaved target dsDNA

Biochemical studies of Cas9 previously revealed that the Cas9-guide RNA complex remains bound to both ends of the double-strand DNA break after cleavage (Sternberg et al., 2014). This is likely a consequence of extensive interactions with both the PAM-proximal and PAM-distal ends of the cleaved dsDNA target. In contrast to Cas9, Cas12a cleaves its target DNA distal from the PAM. Consequently, Cas12a engages in very few interactions with the PAM-distal cleavage product of a target dsDNA substrate. To determine the fate of target DNA after Cas12a-mediated cleavage, we utilized dsDNA targets in which one of the four termini (*i.e.* either the 5’-TS, 3’-TS, 5’-NTS, or 3’-NTS) was labeled with a covalently attached fluorophore. All four target DNAs were efficiently bound by the catalytically inactive FnCas12a^E1006Q-R1218A^-crRNA complex, as judged by fluorescence-detection size exclusion chromatography (**Figure S6**). Upon incubation with a catalytically active FnCas12a-crRNA complex, the target DNAs were cleaved (**Figure 4**). The PAM-proximal cleavage product (*i.e.* 3’-TS and 5’-NTS labeled DNA) eluted together with the FnCas12a-crRNA complex, indicating that the PAM-proximal DNA remains bound to FnCas12a after cleavage. In contrast, PAM-distal DNA (i.e. 5’-TS and 3’-NTS labeled DNA) did not co-elute with the FnCas12a-crRNA complex, indicating that the PAM-distal DNA is released from the FnCas12a-crRNA complex upon cleavage. These findings are in agreement with a recent single molecule fluorescence imaging study (Singh et al., 2018) and with a recent crystal structure of FnCas12a bound to a cleaved DNA target (Stella et al., 2017), in which the REC2 domain is also moved away from the RuvC catalytic site. This suggests that the Cas12a-crRNA complex maintains a catalytically active conformation after dsDNA cleavage. Moreover, release of the PAM-distal DNA product might be essential to facilitate dsDNA-activated *trans*-acting deoxyribonuclease activity as it clears the catalytic site.

**Figure 4.**
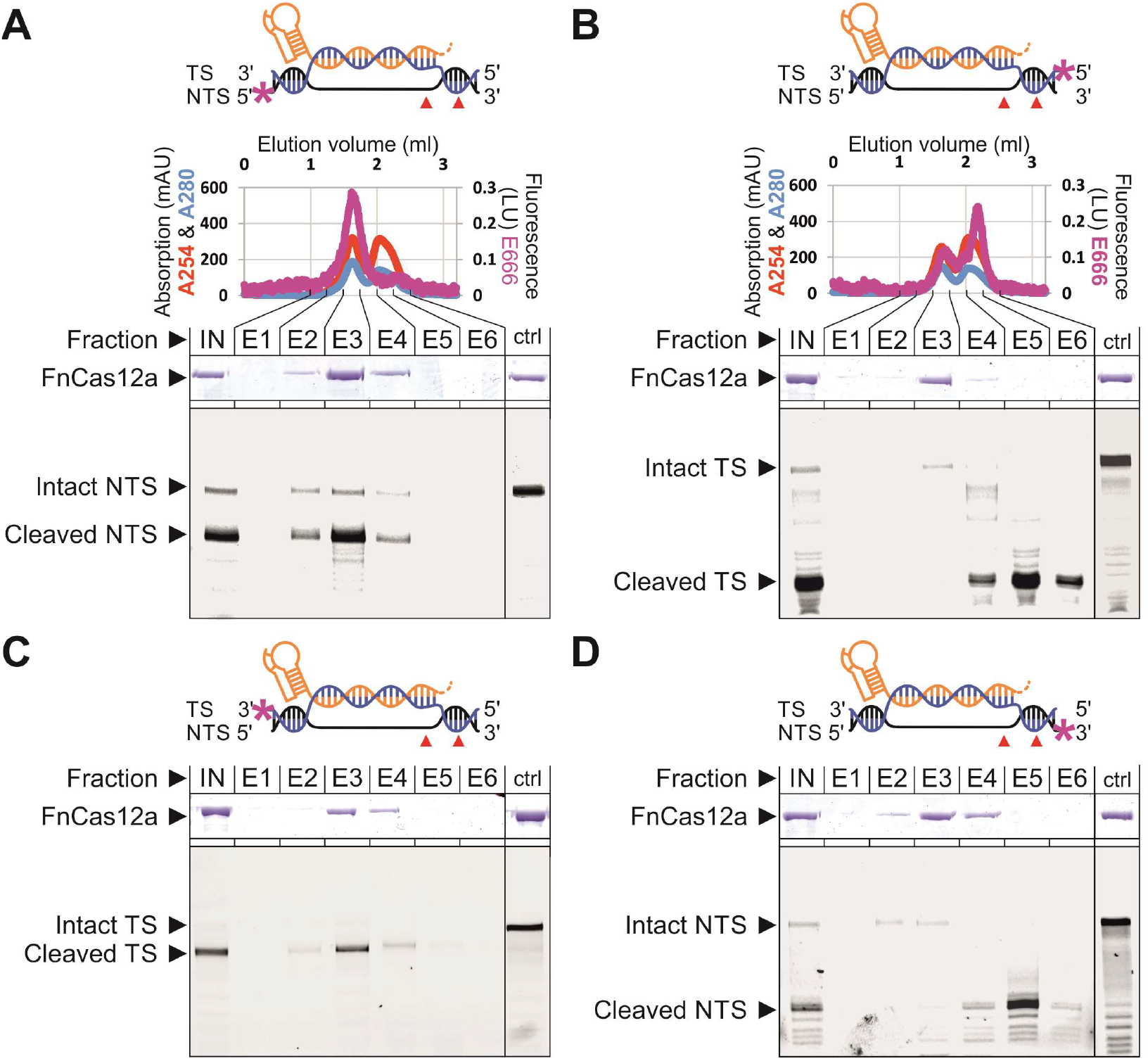
Cas12a-crRNA releases PAM-distal DNA and remains bound to the PAM-proximal cleavage product. (A-D) FnCas12a-crRNA complexes were incubated with fluorophore-labeled dsDNA targets and analyzed by fluorescence-detection size exclusion chromatography. Elution fractions were further analyzed for protein and fluorophore-labeled nucleic acid content by SDS-PAGE and 20% denaturing (7 M Urea) polyacrylamide gel electrophoresis, respectively. IN: HPLC input; E1-6: Elution fractions; ctrl: Control sample containing catalytic mutant FnCas12a^E1006Q-R1218A^ instead of FnCas12a. All HPLC chromatograms and control HPLC runs (containing no protein or FnCas12a^E1006Q-R1218A^) are shown in **Figures S6** and **S7**.

## DISCUSSION

Recent studies have shown that Cas12a enzymes exhibit two DNase activity modes (Chen et al., 2018; Li et al., 2018). Besides *cis*-cleavage of complementary DNA targets, Cas12-family nucleases can catalyze cleavage of non-complementary ssDNA substrates in *trans* upon activation by a complementary DNA target. The target DNA-induced, *trans*-acting DNAse activity has been exploited to develop sensitive technologies for nucleic acid detection (Chen et al., 2018; Gootenberg et al., 2018). By combining insights from two novel structures of FnCas12a-crRNA-DNA complexes and corroborating biochemical experiments, this study provides a revised model for both *cis*- and *trans*-acting deoxyribonuclease activities of Cas12a (**Figure 5**). Hybridization of Cas12a-bound crRNA and TS DNA results in allosteric activation of the RuvC catalytic site, turning Cas12a into a non-specific ssDNase. While this activity can also be activated by dsDNA binding, this requires the presence of a cognate PAM in the dsDNA target. Based on analogies with Cas9, we hypothesize that PAM recognition is essential to initiate local TS-NTS melting to allow base pairing of the TS to the seed segment of crRNA, followed by processive crRNA-TS DNA hybridization and simultaneous TS-NTS unwinding. R-loop formation subsequently results in ordered sequential cleavage of the NTS and the TS. The existence of a conserved NTS binding groove in the RuvC domain, together with the observed dependence of TS cleavage on prior NTS cleavage, collectively suggest that the NTS initially occupies the RuvC catalytic site, thereby blocking TS from accessing the RuvC catalytic site. Furthermore, NTS cleavage might facilitate local melting of the PAM-distal TS-NTS duplex, which is essential to position the TS in the RuvC catalytic site in a single-stranded form. Based on similarities with crystal structures of the related Cas12b nucleases, we assume that the Cas12a Nuc domain induces the TS to adopt a kinked conformation, which promotes its binding in the RuvC site. Upon sequential cleavage of the two DNA strands of a dsDNA target, Cas12a remains in a catalytically competent state, with its RuvC active site exposed to solvent and poised to catalyze cleavage of non-cognate ssDNA substrates in *cis*. However, the exact mechanism by which ssDNA substrates bind in and are cleaved by the RuvC catalytic site remains to be determined.

**Figure 5.**
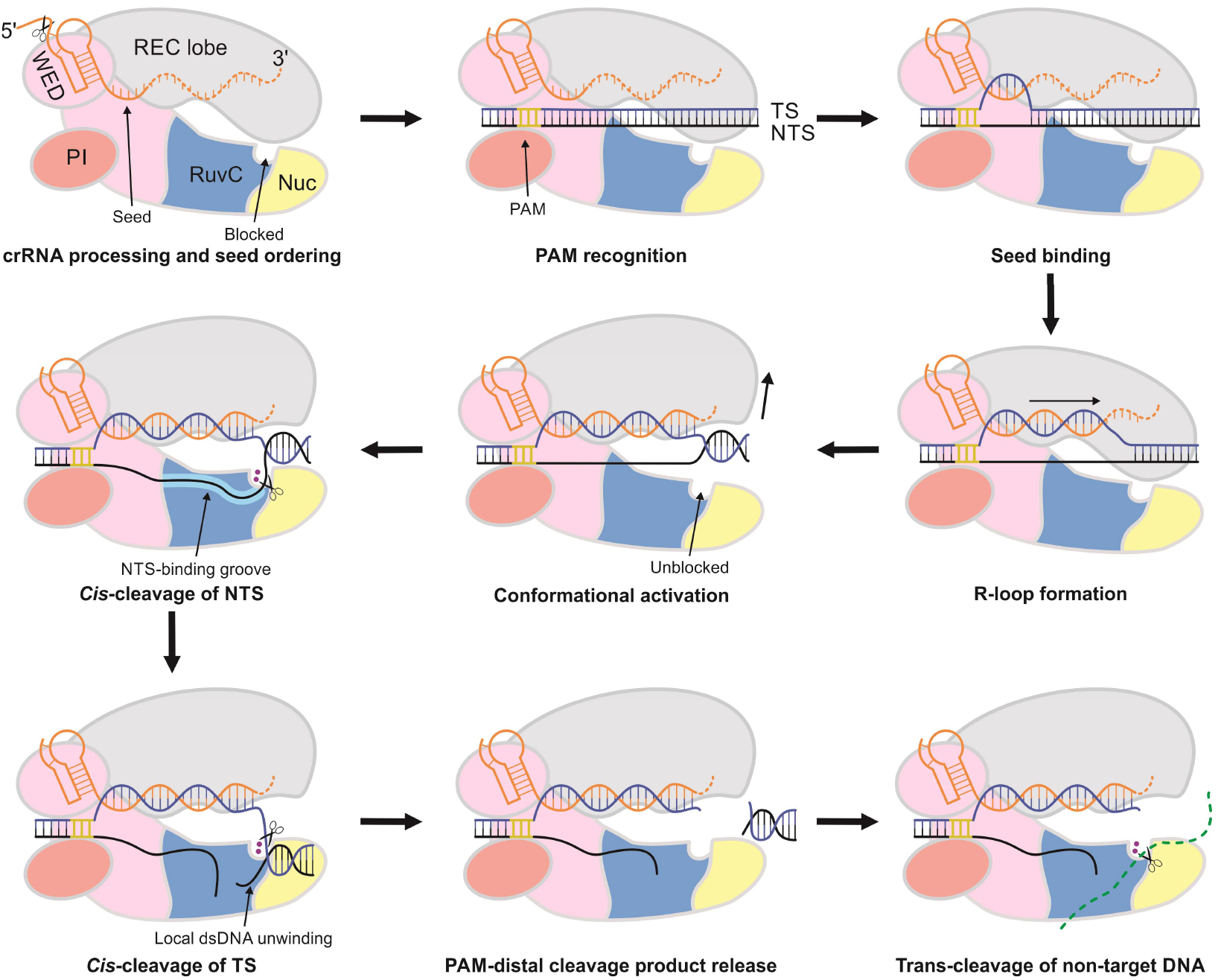
Schematic model of crRNA-guided target DNA binding and *cis*- and *trans*-cleavage of DNA by Cas12a. Cas12a processes its own crRNA guide and pre-orders nucleotides 1-5 in of the crRNA seed segment. PAM recognition by the WED and PI domains promotes target DNA unwinding and places the DNA target strand in register with the pre-ordered crRNA seed sequence. The R-loop is formed by processive crRNA-TS hybridization and simultaneous TS-NTS unwinding. crRNA-TS heteroduplex formation induces conformational changes in the REC lobe resulting in allosteric unblocking of the catalytic site in the RuvC domain. The NTS binding groove guides the displaced NTS toward the catalytic site, resulting in NTS cleavage. Subsequently, further unwinding of the PAM-distal TS-NTS duplex allows placement of the TS into the catalytic site, resulting in TS cleavage. The PAM-distal dsDNA is released, while the PAM-proximal dsDNA remains bound to the Cas12a-crRNA complex. Cas12a remains in a catalytically activated conformation, which allows for cleavage of non-target DNAs in *trans*.

The insights provided by this study likely apply to other Cas12a orthologs as well as to Cas12b nucleases. Similar to Cas12a, Cas12b demonstrates *cis*-acting and *trans*-acting deoxyribonuclease activities (Chen et al., 2018). Moreover, the Cas12b RuvC nuclease domain is responsible for cleavage of both the TS and NTS DNA strands in a dsDNA target (Liu et al., 2017; Yang et al., 2016). These findings are important in the context of the biological function of type V CRISPR-Cas systems in genome defense as they imply that these systems can target both ssDNA and dsDNA invasive nucleic acid elements. Furthermore, following cleavage of complementary dsDNA, the persistence of *trans*-acting DNase activity might contribute to the interference mechanism and invader clearance.

Finally, these insights have implications for further development and utilization of Cas12a as a genome editing tool. FnCas12a and other Cas12a orthologs have successfully been used for genome editing in bacteria, yeast, insects, plants, and mammalian cells, including human cell lines (Swarts and Jinek, 2018). Cas12a does not only cleave target genomic dsDNA in *cis*, but could also cleave non-target ssDNA in *trans*, thus potentially targeting transcription or replication intermediates, or ssDNA templates that are used for homology-directed repair (Richardson et al., 2016). Furthermore, the asymmetric retention and release of DSB ends upon Cas12a-catalyzed cleavage suggests that editing efficiency or DNA repair outcomes might be dependent on the orientation of the bound Cas12a, particularly in the context of multiplexed genome editing. The significance of *trans*-acting DNase activity for genome editing awaits further investigation. In conclusion, our studies revealed critical insights into the molecular mechanisms of target DNA binding, cleavage, and release by type V CRISPR effector nucleases, which might have important implications for genome editing.

## Methods

### FnCas12a expression and purification

FnCas12a and FnCas12a mutants were expressed and purified as described previously (Swarts et al., 2017). A detailed step-by-step protocol can be found online at bio-protocol (https://bio-protocol.org/e2842).

### Crystallization and structure determination

The ternary FnCas12a-crRNA-TS complex was reconstituted by combining purified apo-FnCas12a^E1006Q^ in SEC buffer (500 mM KCl, 20 mM HEPES (pH 7.5), 1 mM DTT) at a concentration of 10 mg.ml^−1^ with synthetic crRNA1 (obtained from Integrated DNA Technologies, Table S1) in a 1:1.4 molar ratio (FnCas12a:crRNA1) in the presence of 5 mM MgCl_2_ and incubating for 10 min at RT. Next, the TS DNA oDS311 (Table S1) was added in a 1:14:1.6 ratio (FnCas12a:crRNA1:oDS311). The complex was crystallized at 20°C using the hanging drop vapor diffusion method by mixing equal volumes of protein and reservoir solution. Initial spherulites were obtained at a protein concentration of 8.3 mg.ml^−1^ with 1.6 M trisodium citrate as reservoir solution. Crystal growth was optimized by iterative microseeding. Data was collected from a crystal that was grown for four weeks at 20 °C in a protein concentration of 8.3 mg.ml^−1^ and 1.6 M trisodium citrate as reservoir solution. Crystals were transferred to a cryoprotectant solution (1.44 M trisodium citrate, 10% ethylene glycol) and flash-cooled in liquid nitrogen.

The ternary FnCas12a-crRNA-DNA complex was reconstituted and crystallized as described previously for the FnCas12a-crRNAX-DNA ternary complex (Swarts et al., 2017). Diffraction data was obtained using a crystal grown for four months at 20°C in refinement screens with a final protein concentration of 7.7 mg.ml^−1^ and a reservoir solution containing 0.1 M Bis-Tris Propane (BTP; pH 6.5), 0.2 M KSCN, and 17.5 % polyethylene glycol 3,400. Crystals were transferred to a cryoprotectant solution (0.1 M BTP (pH 6.5), 0.2 M KSCN, 20% polyethylene glycol 3,400, 15% (v/v) ethylene glycol, and 5 mM MgCl_2_ and flash-cooled in liquid nitrogen.

X-ray diffraction data were measured at beamline X06DA (PXIII) of the Swiss Light Source (Paul Scherrer Institute, Villigen, Switzerland). Data were indexed, integrated, and scaled using XDS. Crystals of the FnCas12a-crRNA-TS ternary complex diffracted to a resolution of 2.8 Å and belonged to space group *P*2_1_2_1_2_1_, with two copies of the complex in the asymmetric unit. The structure of the FnCas12a-crRNA-TS complex was solved by molecular replacement in phenix.phaser using a modified FnCas12a-crRNAX-dsDNA (PDB: 5NFV) as search model. Phases obtained using the initial molecular replacement solution were improved by density modification using phenix.resolve (Terwilliger and Terwilliger, 2004) and phenix.morph_model (Terwilliger et al., 2013). The atomic model was built manually in Coot (Emsley and Lohkamp, 2010) and refined using phenix.refine (Afonine et al., 2012). The final binary complex model contains crRNA1 residues (−18)–(+21), target strand residues (−20)−(−1) and FnCas12a^E1006Q^ residues 1-1299, except for residues 550-553, 1009-1013, 1156-1162, and 1277-1279, which lack ordered electron density.

The crystal of the FnCas12a-crRNA-dsDNA ternary complex diffracted to a resolution of 2.65 Å and belonged to space group *P*2_1_, with one copy per asymmetric unit. The structure was solved by molecular replacement in phenix.phaser using a modified FnCas12a-crRNAX-dsDNA (PDB: 5NFV) as search model. The atomic model was built manually in Coot and refined using phenix.refine. The final ternary complex model contains crRNA residues (−18)–20, target DNA strand nucleotides (−27)–10, and non-target strand nucleotides (−8*)-13* and 19*-29*, and FnCas12a residues 1–1300, except for residues 850-851, 1135-1136, and 1153-1166, which lack ordered electron density.

### Structure analysis

Root Means Square Deviations (rmsd) of structure alignments were calculated using Coot LSQ superpose (Emsley and Lohkamp, 2010). Domain movements were analysed using DynDom (Hayward and Lee, 2002). Intramolecular interactions were analysed using PDBePISA (Krissinel and Henrick, 2007).

### *Trans*-cleavage experiments

To generate dsDNA targets, TS and NTS oligonucleotides (100 μM, oDS329-oDS342, Table S1) were mixed in a 1:4 molar ratio (TS:NTS). The strands were annealed by incubation at 95°C for 5 min, followed by slow cooling to room temperature. The samples were further diluted in H_2_O to a concentration of 10 μM. For *trans*-cleavage experiments, FnCas12a and FnCas12^E1006Q^ (10 μM in SEC buffer) were mixed with crRNA1 (10 μM in H_2_O) in the presence of 5 mM MgCl_2_ and incubated at 37°C for 10 min to allow binary complex formation. Target ssDNAs oDS302 or oDS311 (Table S1, 10 μM in H_2_O) or target dsDNAs (10 μM in H_2_O) and M13 circular ssDNA or dsDNA substrates (Table S2, 100 ng.μl^−1^, New England Biolabs) were added. The final reaction (20 μl) contained final concentrations of 0.5x SEC buffer, 2.5 μM FnCas12a or FnCas12a^E1006Q^, 1.5 μM crRNA1, 5 mM MgCl_2_, 1.5 μM trans-activating ssDNA or dsDNA, and 10 ng.μl^−1^ circular DNA substrate. Samples were incubated for 90 min at 37 °C. Reactions were stopped by adding EDTA and Proteinase K (Thermo Fisher Scientific) to final concentrations of 80 mM and 0.8 mg.ml^−1^, respectively, and incubating for 30 min at 37°C. Trans-activating dsDNA was removed from the M13 circular ssDNA using the NucleoSpin® Gel and PCR Clean-up kit (MACHERY-NAGEL) according to the instructions of the manufacturer, with 7-fold excess of NTI DNA binding buffer. 6x DNA loading dye (Thermo Fisher Scientific) was added to the sample and the DNA was resolved on 0.8% agarose gels stained with GelRed Nucleic Acid Gel Stain (Biotum) and visualized using a ChemiDoc Touch gel imager (Bio-Rad).

### Order of strand cleavage experiments

The dsDNA targets with modified backbones were generated by mixing TS and NTS oligonucleotides (100 μM, Table S1) in a 1:2 ratio (unlabeled:Cy5-labeled). The strands were annealed by incubation at 95°C for 5 min, followed by slow cooling to room temperature. The samples were mixed in a 1:1 ratio with native loading dye (0.5x SEC buffer, 5mM MgCl2, 50% glycerol) and resolved on a 10% native polyacrylamide gel. Gel regions containing fluorophore-labeled dsDNA were identified using a Typhoon FLA 9500 gel imager (GE Healthcare) and cut out. The gel was crushed and soaked in H_2_O for two days before the supernatant was extracted. The dsDNA was isolated from the supernatant by ethanol precipitation and dissolved in H_2_O. The final concentration of the dsDNA targets was ~0.3 μM.

FnCas12a or FnCas12a^E1006Q-R1218A^ (10 μM in SEC buffer), crRNA-λ (10 μM), and MgCl_2_ (50 mM) were mixed and incubated for 10 min at 37°C before the target dsDNA (0.3 μM) was added. The final 20 μl reaction contained final concentrations of 0.25x SEC buffer, 2.5 μM FnCas12a or FnCas12a^E1006Q-R1218A^, 1 μM crRNA-μ, 2.5 mM MgCl_2_, and 30 nM target dsDNA, and was incubated for 30 min at 20°C. Reactions were stopped by adding EDTA and Proteinase K (Thermo Fisher Scientific) to final concentrations of 80 mM and 0.8 mg.ml^−1^, respectively, and incubating for 30 min at 37°C. Samples were mixed 1:1 with 2x loading dye (5% glycerol, 90% formamide, and 2.5 mM EDTA), heated for 10 min at 95°C, and resolved on a 20% denaturing (7 M Urea) polyacrylamide gel. Fluorescence of the Cy5-labeled DNA was detected using a Typhoon FLA 9500 gel imager (GE Healthcare).

### High-performance liquid chromatography experiments

Labeled and unlabeled target strand oligonucleotides (100 μM, Table S1) and non-target strand oligonucleotides (100 μM, Table S1) were mixed in a 1:2 ratio (labeled:unlabeled) and annealed by incubation at 95°C for 5 min, followed by slow cooling to room temperature. The formed dsDNA targets were diluted in water to 10 μM. Prior to high-performance liquid chromatography (HPLC) analysis, FnCas12a or FnCas12a^E1006Q-R1218A^, crRNA-λ (10 μM), MgCl_2_ (50 mM), SEC buffer, and target DNA (10 μM) were mixed. The resulting 100 μl reaction contained final concentrations of 0.5x SEC buffer, 2 μM FnCas12a or FnCas12a^E1006Q-R1218A^, 1 μM crRNA-λ, 2.5 mM MgCl_2_, and 0.25 μM target dsDNA, and was incubated for 1h at 37°C. Samples were resolved by HPLC on a Superdex 200 5/150 size exclusion column (GE Life Science) with a 3-ml bed volume and 20 bar pressure limit, which was equilibrated and run in 0.5x SEC at a flow rate of 0.1 ml min^−1^. Absorption at 280nm and 260nm and the fluorescent signal of Cy5- or ATTO532-labeled DNA were recorded throughout the run. For each target dsDNA, fractions were collected and subsequently analysed by SDS-PAGE gel electrophoresis for protein content and by 20% denaturing (7 M Urea) PAGE for DNA content. Fluorescence of the Cy5- or ATTO532-labeled DNA was detected using a Typhoon FLA 9500 gel imager (GE Healthcare).

### Data and Software Availability

The atomic coordinates and structure factors reported in this paper will be deposited in the Protein Data Bank (PDB). The unprocessed image files used to prepare the figures in this manuscript will be deposited in Mendeley Data.

## Author contributions

D.C.S. and M.J. designed experiments. D.C.S. prepared and crystallized the Cas12a complexes, collected X-ray data, determined crystal structures and carried out biochemical assays. D.C.S. and M.J. wrote the manuscript.

## Acknowledgements

We are grateful to Meitian Wang, Vincent Olieric, and Takashi Tomizaki at the Swiss Light Source (Paul Scherrer Institute, Villigen, Switzerland) for assistance with X-ray diffraction measurements. We thank members of the Jinek group for discussions and critical reading of the manuscript. This work was supported by a Swiss National Science Foundation (SNSF) Project Grant to M.J. (SNSF 31003A_149393) and by long-term postdoctoral fellowships from the European Molecular Biology Organization (EMBO) to D.C.S (ALTF 179-2015 and aALTF 509-2017). M.J. is International Research Scholar of the Howard Hughes Medical Institute and Vallee Scholar of the Bert L & N Kuggie Vallee Foundation.

---

**Figure S1.**
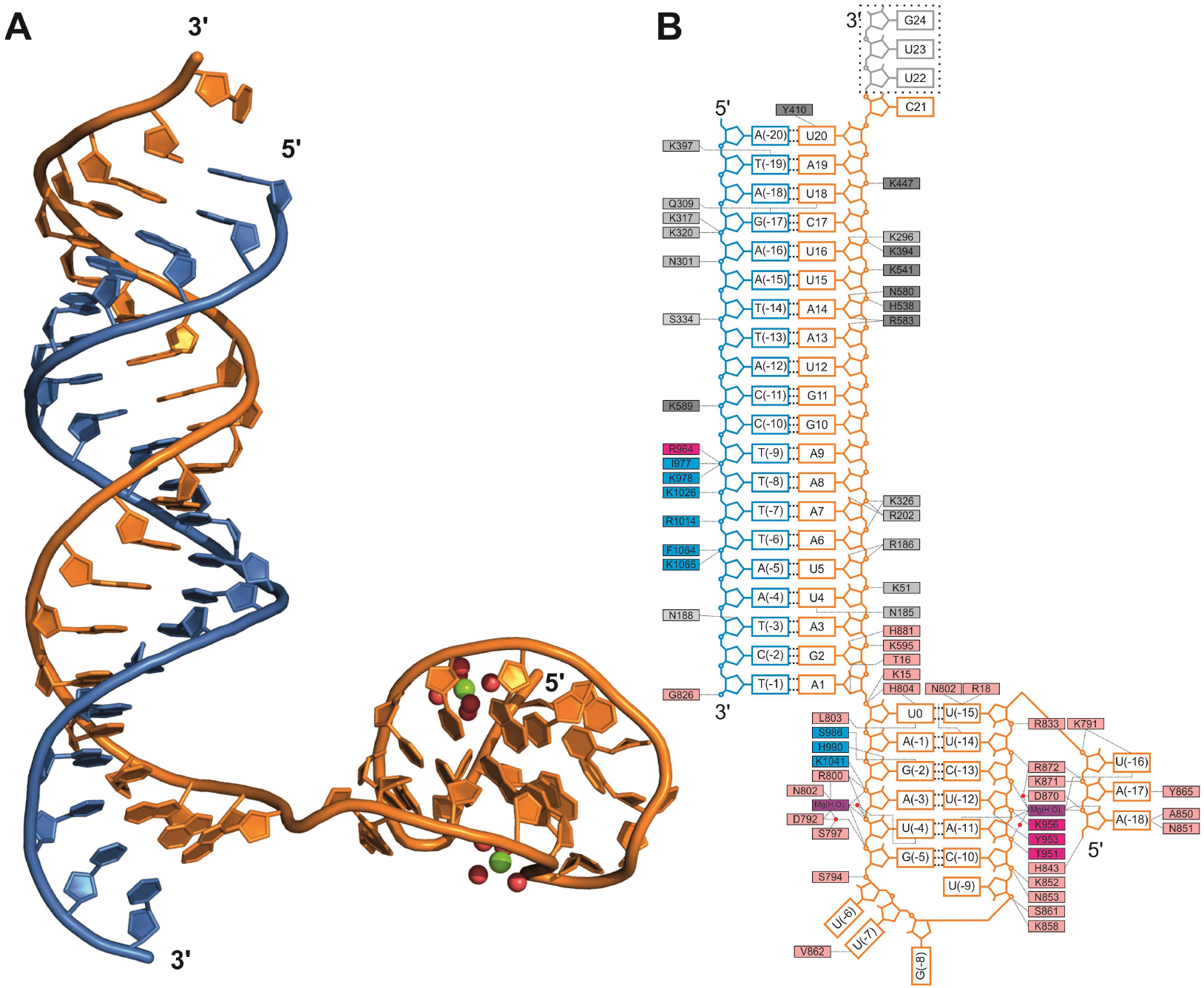
Details of FnCas12a-crRNA-TS interactions. Related to Figure 1 and 2. (A) Structure of nucleic acids in the crystal structure of FnCas12a-crRNA complex bound to a target strand. Water molecules are depicted as red spheres; Mg^2+^ ions are depicted as green spheres. (B) Schematic representation of hydrogen bonding interactions between FnCas12a, nucleic acids, and divalent cations in the ternary structure. FnCas12a residues are colored according to their domains (see Figure 1A). Nucleotides colored grey are not ordered in the structure. Base pairs are indicated with thick dashed lines, while other hydrogen bonds are indicated with thin dashed lines. Red circles indicate water-mediated hydrogen bonding. Intra-crRNA pseudoknot hydrogen bonds in the crRNA are not displayed for clarity.

**Figure S2.**
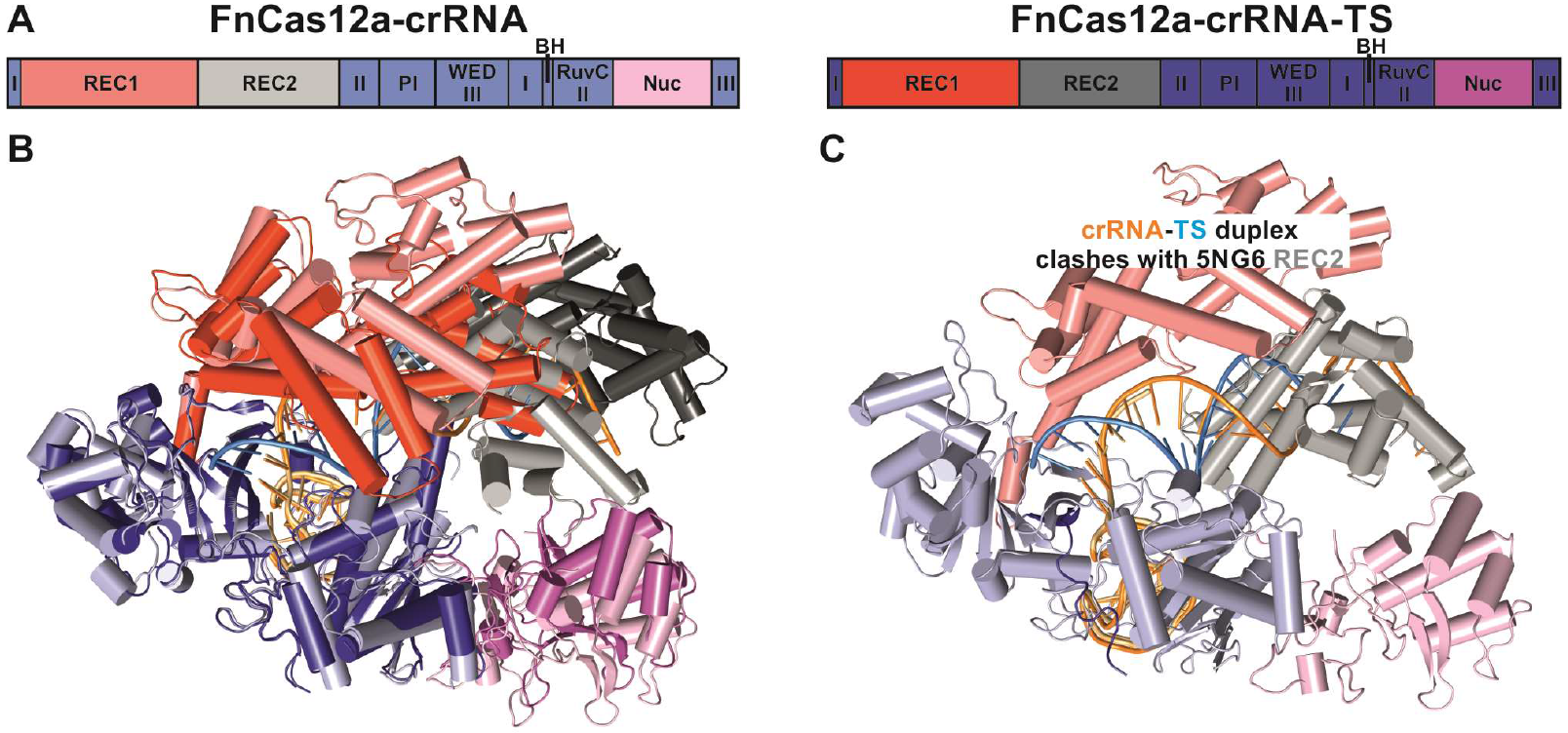
crRNA-TS hybridization induces conformational changes in FnCas12a. Related to Figure 1 and 2. (A) Schematic diagram of the domain organization and coloring of the FnCas12a complexes in panel B and C. (B) Alignment of the binary FnCas12a-crRNA complex (PDB: 5NG6) and the ternary Fncas12a-crRNA-TS complex structures. (C) The crRNA-TS duplex of the ternary FnCas12a-crRNA-TS complex cannot be accommodated by the binary FnCas12a-crRNA complex due to steric clashes with the REC2 domain.

**Figure S3.**
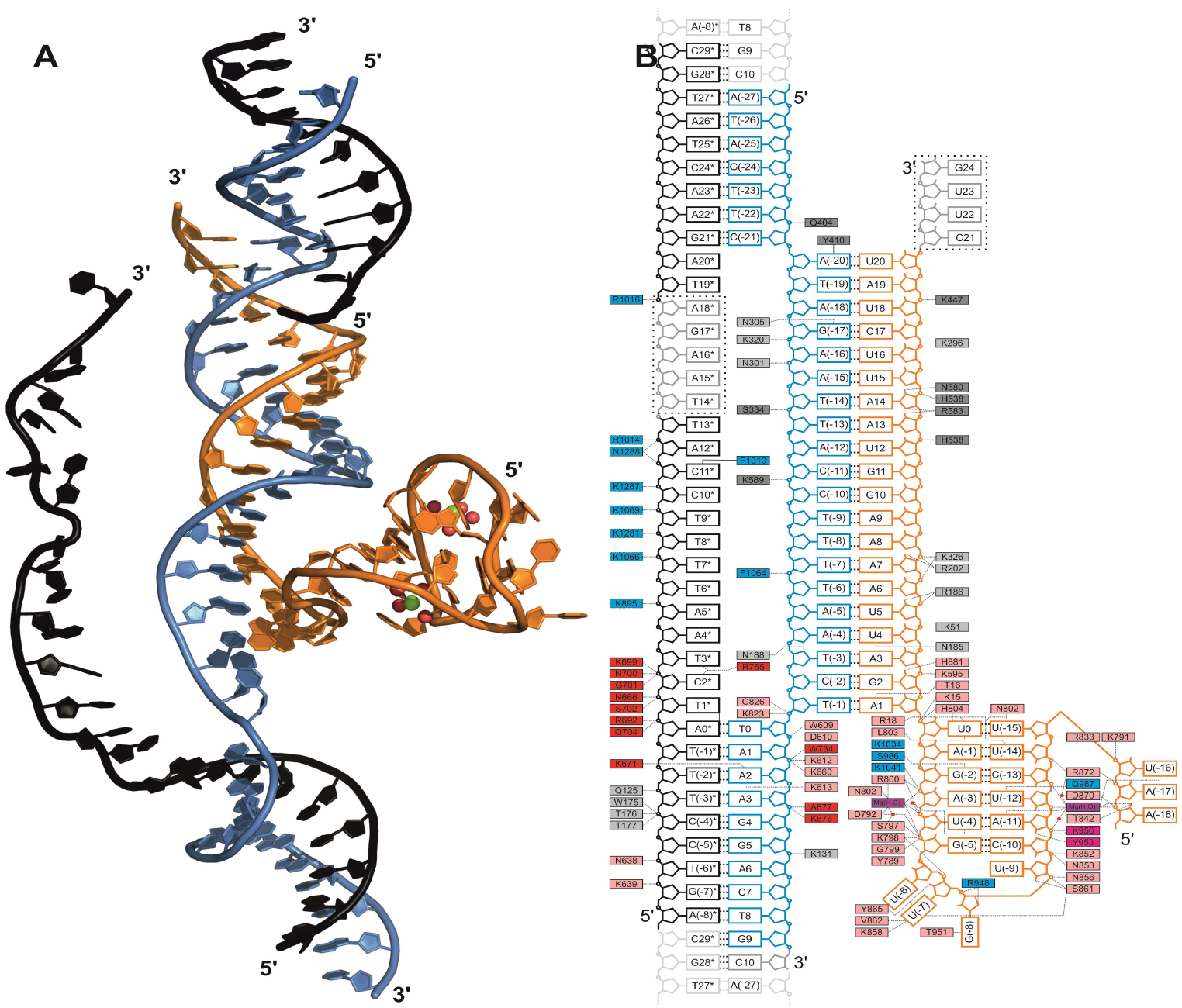
Details of FnCas12a-crRNA-dsDNA interactions. Related to Figure 3. (A) Structure of nucleic acids in the structure of FnCas12a-crRNA complex bound to a dsDNA target. Water molecules are depicted as red spheres; Mg^2+^ ions are depicted as green spheres. (B) Schematic representation of hydrogen bonding interactions between FnCas12a, nucleic acids, and divalent cations in the ternary FnCas12a-crRNA-dsDNA structure. FnCas12a residues are colored according to their domains (see Figure 1A). Nucleotides colored grey are not ordered in the structure. Base pairs are indicated with thick dashed lines, while other hydrogen bonds are indicated with thin dashed lines. Red circles indicate water-mediated hydrogen bonding. Intra-crRNA pseudoknot hydrogen bonds in the crRNA are not displayed for clarity.

**Figure S4.**
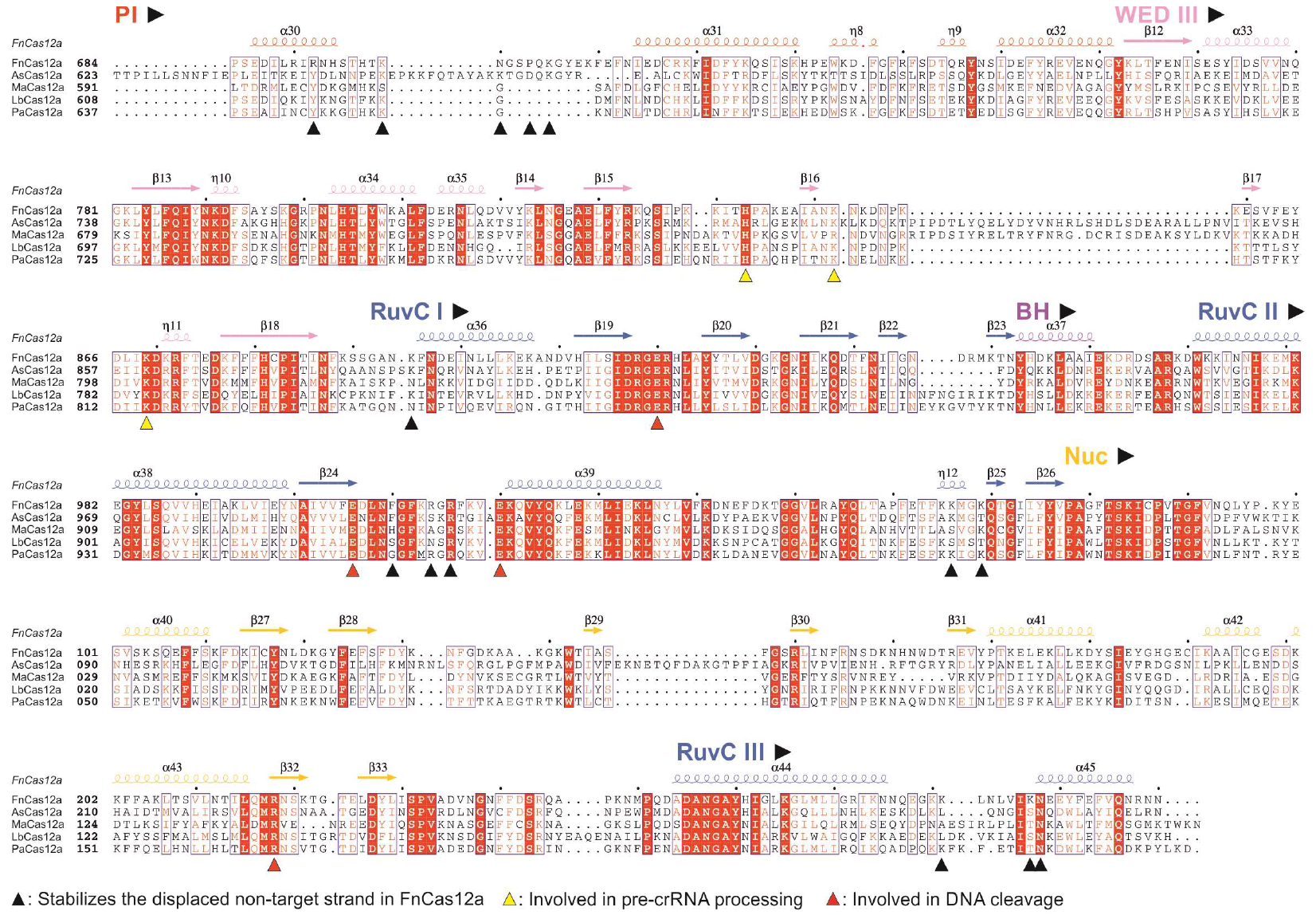
Multiple sequence alignment of FnCas12a orthologs. Related to Figure 3. Clustal Omega (Larkin et al., 2007) was used to generate a Multiple sequence alignment of Cas12a protein sequences of *Francisella novicida* U122 (FnCas12a), *Acidaminococcus sp*. BV3L6 (AsCas12a), *Methanomethylophilus alvus* Mx1201 (MaCas12a), and *Lachnospiraceae bacterium* ND2006 (LbCas12a). The Clustal Omega sequence alignment and the structural information from the structure of the binary FnCas12acrRNA complex were used as input for ESPript 3.0 (http://espript.ibcp.fr) (Robert and Gouet, 2014) to align secondary structure features to the sequence alignment. Residues important for specific FnCas12a functions are indicated with colored triangles.

**Figure S5.**
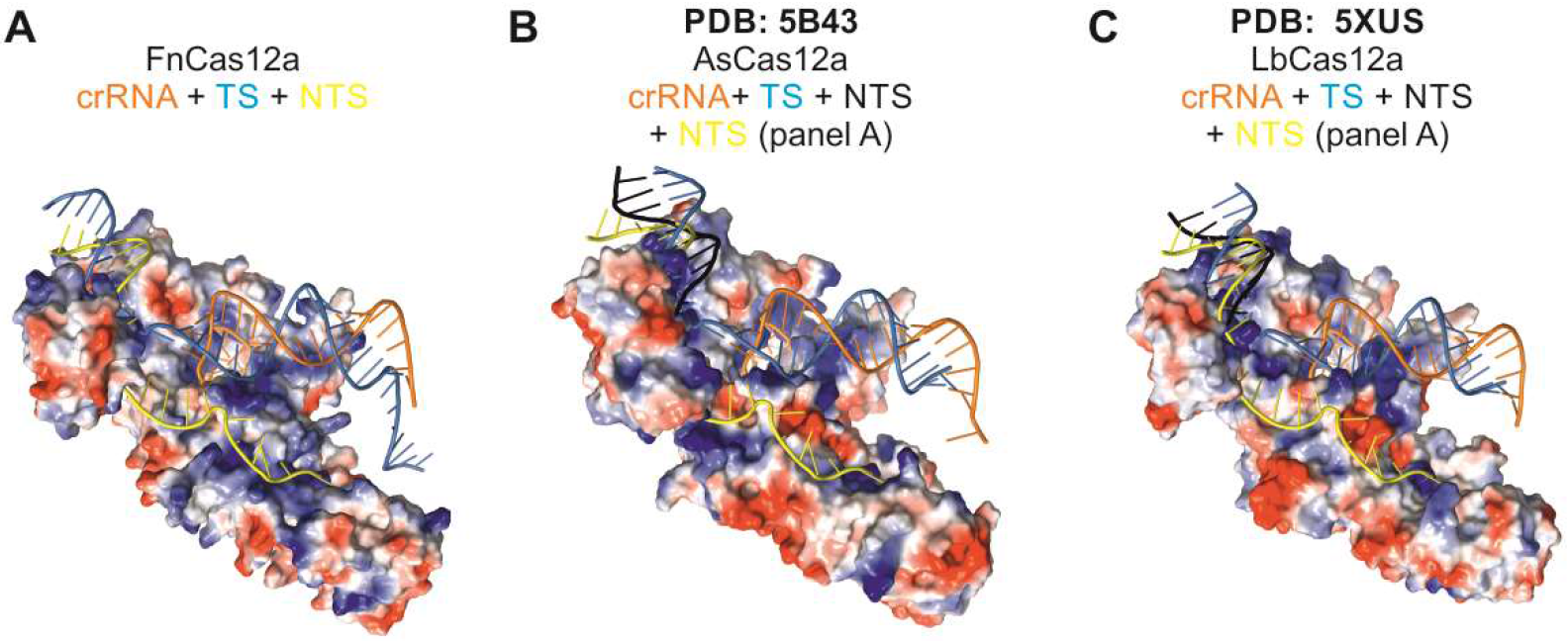
The NTS binding groove is structurally conserved in Cas12a orthologs. Related to Figure 3. (A) Surface electrostatic potential map of the NUC lobe of the FnCas12a-crRNA-dsDNA complex (as in Figure 3 panel B). Blue, positively charged region; red, negatively charged region. The REC lobe is omitted for clarity. (B) Surface electrostatic potential map of the AsCas12a NUC lobe (PDB: 5B43). Blue, positively charged region; red, negatively charged region. The NTS (colored yellow) is modeled based on the structure of R-loop structure of the FnCas12a-crRNA-dsDNA complex. The REC lobe is omitted for clarity. (C) Surface electrostatic potential map of the LbCas12a NUC lobe (PDB: 5XUS). Blue, positively charged region; red, negatively charged region. The NTS (colored yellow) is modeled based on the structure of R-loop structure of the FnCas12a-crRNA-dsDNA complex. The REC lobe is omitted for clarity.

**Figure S6.**
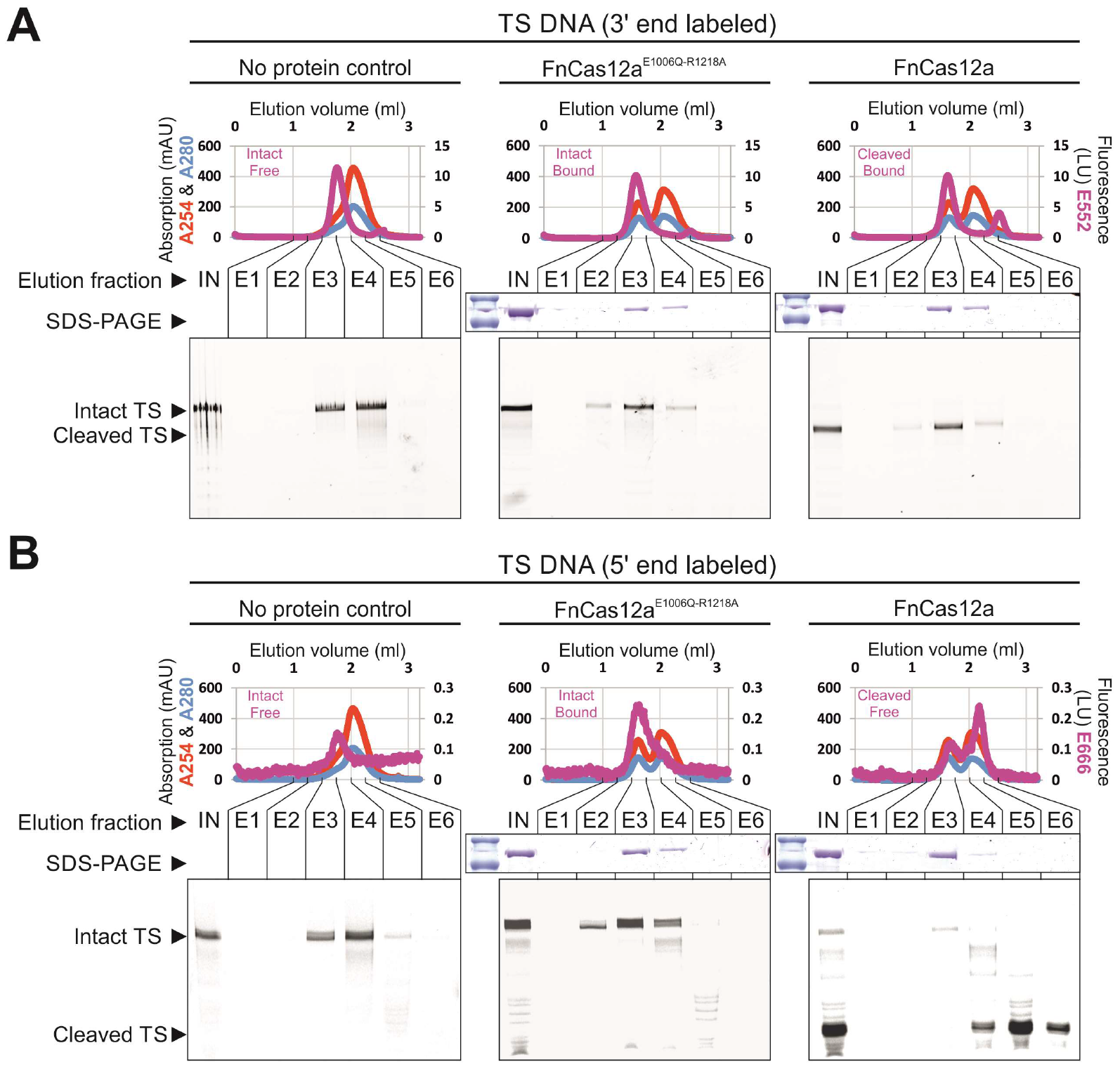
Fate of the target DNA strand after Cas12a-crRNA mediated cleavage. Related to Figure 4. FnCas12a-crRNA or FnCas12a^E1006Q-R1218A^-crRNA complexes were incubated with dsDNA targets containing fluorophore-labeled TS (panel A: TS 3’ end labeled, panel B: TS 5’ end labeled) and analyzed by fluorescence-detection size exclusion chromatography. Elution fractions were further analyzed for protein and fluorophore-labeled nucleic acid content by SDS-PAGE and 20% denaturing (7 M Urea) polyacrylamide gel analyses, respectively. IN: HPLC input; E1-6: Elution fractions; ctrl: Control sample containing catalytic mutant FnCas12a^E1006Q-R1218A^ instead of FnCas12a.

**Figure S7.**
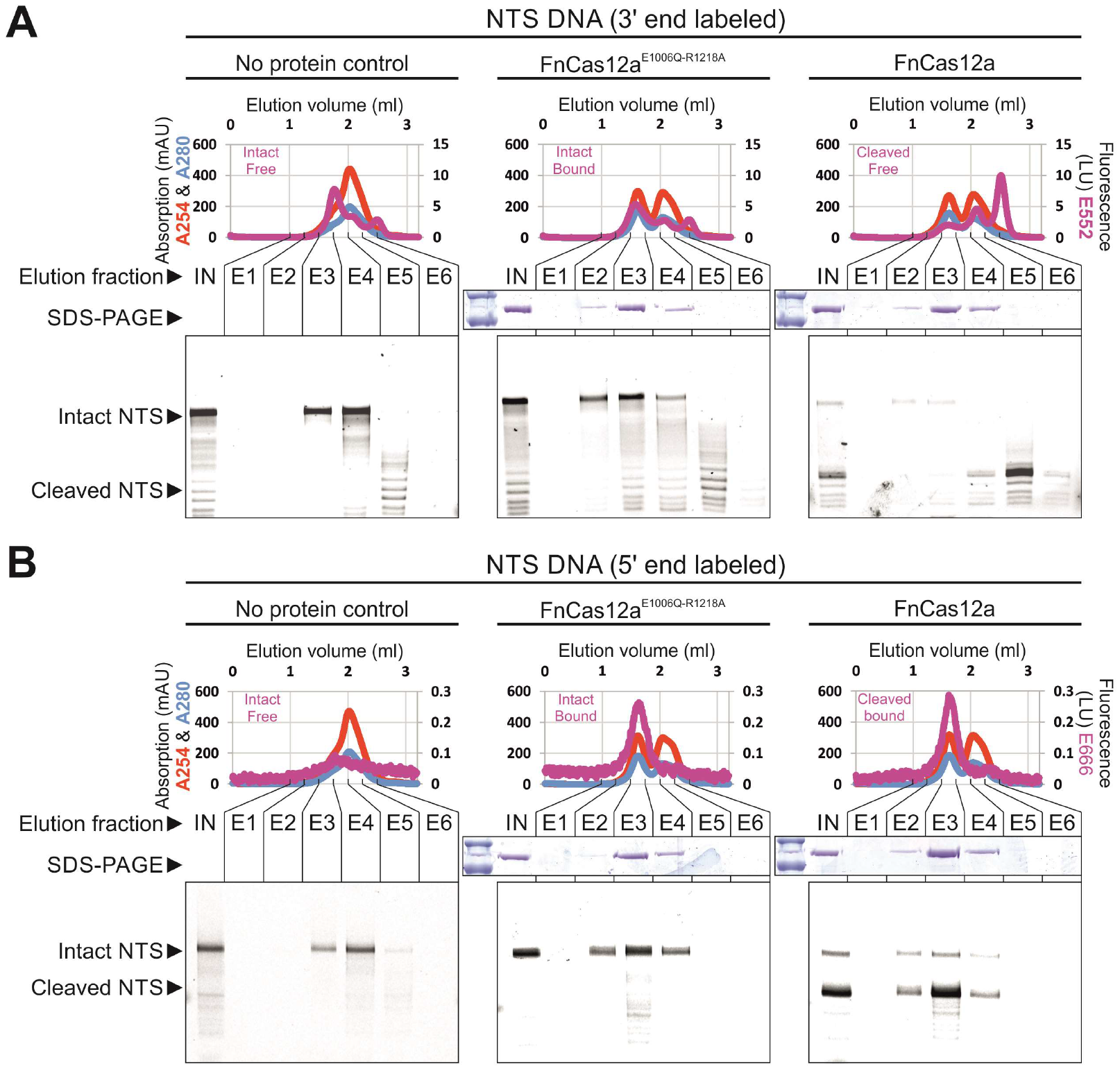
Fate of the non-target DNA strand after Cas12a-crRNA mediated cleavage. Related to Figure 4. FnCas12a-crRNA or FnCas12a^E1006Q-R1218A^-crRNA complexes were incubated with dsDNA targets containing fluorophore-labeled NTS (panel A: NTS 3’ end labeled, panel B: NTS 5’ end labeled) and analyzed by fluorescence-detection size exclusion chromatography. Elution fractions were further analyzed for protein and fluorophore-labeled nucleic acid content by SDS-PAGE and 20% denaturing (7 M Urea) polyacrylamide gel analyses, respectively. IN: HPLC input; E1-6: Elution fractions; ctrl: Control sample containing catalytic mutant FnCas12a^E,006Q-R1218A^ instead of FnCas12a.

**Table S1.**
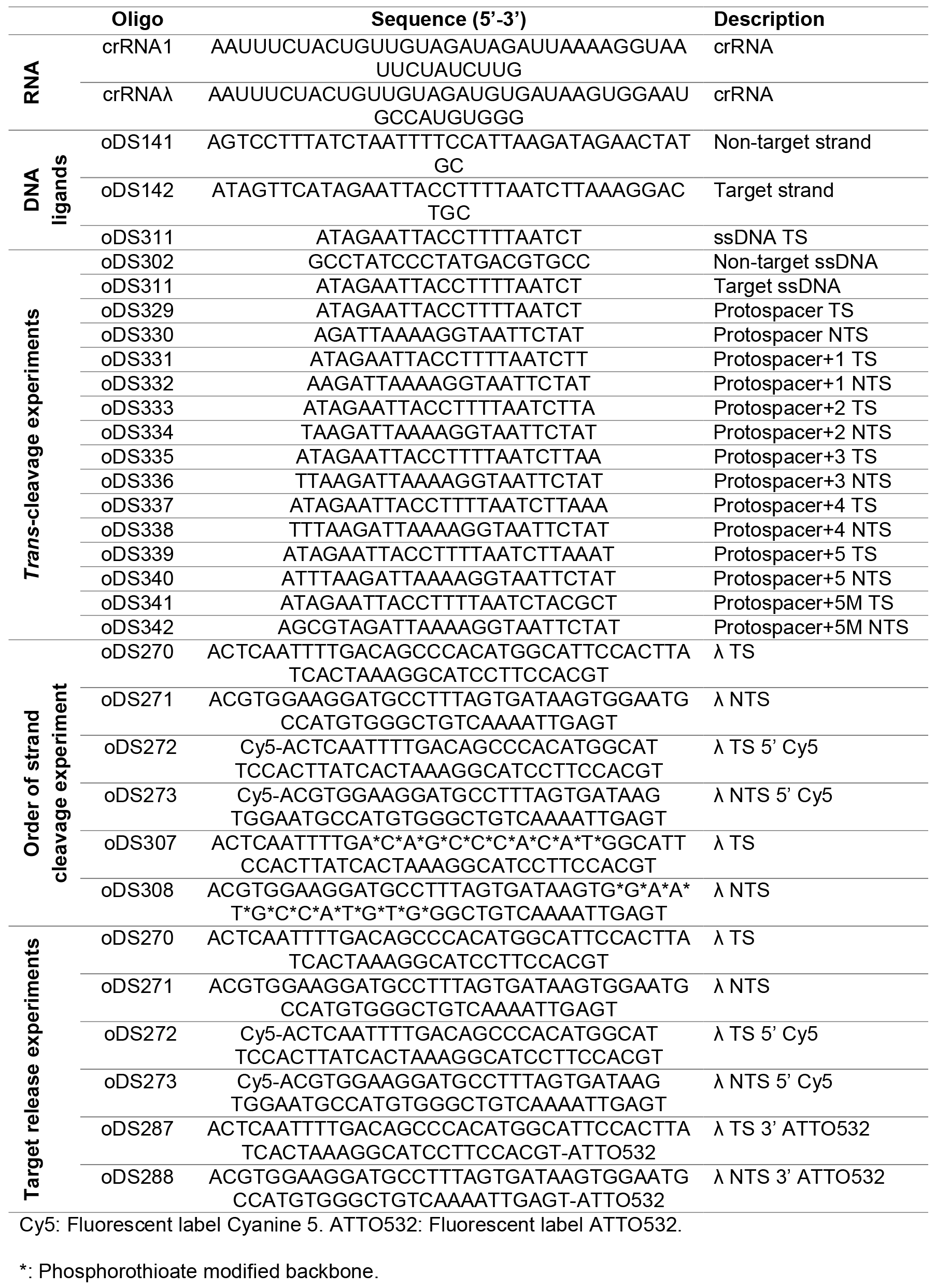
Oligonucleotides used in this study. Related to Figure 1-4

**Table S2.** Plasmids used in this study. Related to Figure 1-4

| **Plasmid** | **Description** | **Source** |
| --- | --- | --- |
| **pDS015** | *F. novicida cas12a* with N-term. His-tag and TEV site in pML-1B. Expression vector for FnCas12a. | (Swarts et al., 2017) |
| **pDS069** | Like pDS015, but with mutation E1006Q | (Swarts et al., 2017) |
| **pDS073** | Like pDS015, but with mutations E1006Q and R1218A | (Swarts et al., 2017) |
| **M13mp18** | Single stranded M13 DNA | New England Biolabs |
| **M13mp18 RF I** | Double stranded M13 DNA | New England Biolabs |

